# Computationally Guided Design of BCR-ABL Tyrosine Kinase Inhibitors

**DOI:** 10.1101/2020.08.20.259887

**Authors:** Chenghao Duan, Simon K. S. Chu, Justin B. Siegel, Jason S. Fell

## Abstract

**BCR-ABL** tyrosine kinase inhibitors (TKI) are used to treat the chronic myeloid leukemia (CML). Many TKI have been developed as the primary treatment to the CML. **Imatinib**, a first generation TKI, directly targets **BCR-ABL** with effective results. As the disease becomes more advanced, patients start to develop resistance to **imatinib**. Due to this effect it is necessary to generate novel treatments for advanced stage CML. Computational tools can predict new drug candidates to target **BCR-ABL**. We have designed two new drug candidates with different levels of modification, based on the predicted structure activity relationships with **BCR-ABL**. These new drug candidates are predicted to have better binding affinities with **BCR-ABL** than **imatinib**, which can be more potent treatments of the disease.

## INTRODUCTION

Chronic myeloid leukemia (CML) is a fatal blood cancer with 1 to 2 cases out of 100,000 people per year and is relatively common in elderly people with a median age of diagnosis of around 65 years old.^1,2^ Nearly 20% to 30% of all CML patients will die within 2 years of diagnosis, and nearly 25% will die each year after that.^3,4,5^ The disease manifests from the fusion of chromosomes 9 and 22, creating what is known as the Philadelphia chromosome, within hematopoietic stem cells.^1^ From this chromosomal fusion the **ABL** tyrosine kinase and the **BCR** genes fuse together to form a unique chimeric **BCR-ABL** oncoprotein. Since the pathogenesis of the disease is stringently related to **BCR-ABL**, primary treatments have been designed to inhibit this enzyme. **BCR-ABL** tyrosine kinase inhibitors (TKI) are a drug family that were developed to bind to the catalytic cleft of **BCR-ABL** to prevent enzyme action.

The first line of treatment of CML is **imatinib** (Gleevec, STI-571; Scheme 1a), and most patients who are treated with imatinib in the diseases early stages achieve durable complete cytogenetic responses long-term event-free survival, and transformation-free survival.^6^ Before **imatinib** (circa 2001) CML patients were treated with combinations of chemotherapy (primarily busulfan) and interferon, which did not prolong patient life and gave an estimated 8-year survival rate of nearly 6%. Since 2001, treatments with **imatinib** have substantially increased the 8-year survival rate of CML patients to 87%.^7^ **Imatinib** binds to the activation loop of **BCR-ABL** which places the enzyme in an inactive conformation.^6^ **Imatinib** contains a pyridine ring, which mimics ATP binding, and a piperazine moiety that binds hydrophobically to the **BCR-ABL** activation loop, preventing enzyme activation, this interaction is showing in figure 1.^8^

**Figure 1.**
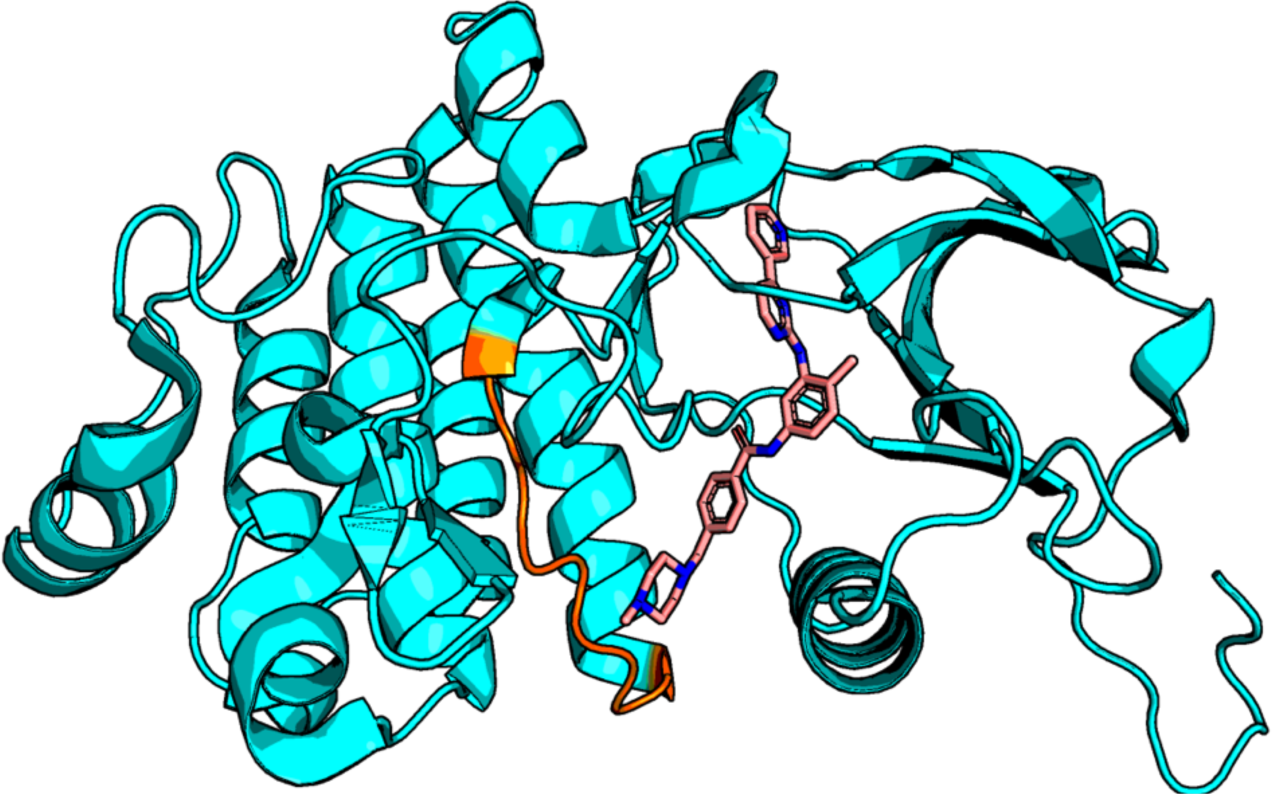
**Imatinib** (pink) within the **BCR-ABL** (teal) active site interacting with the activation loop (orange).

**Scheme 1.**
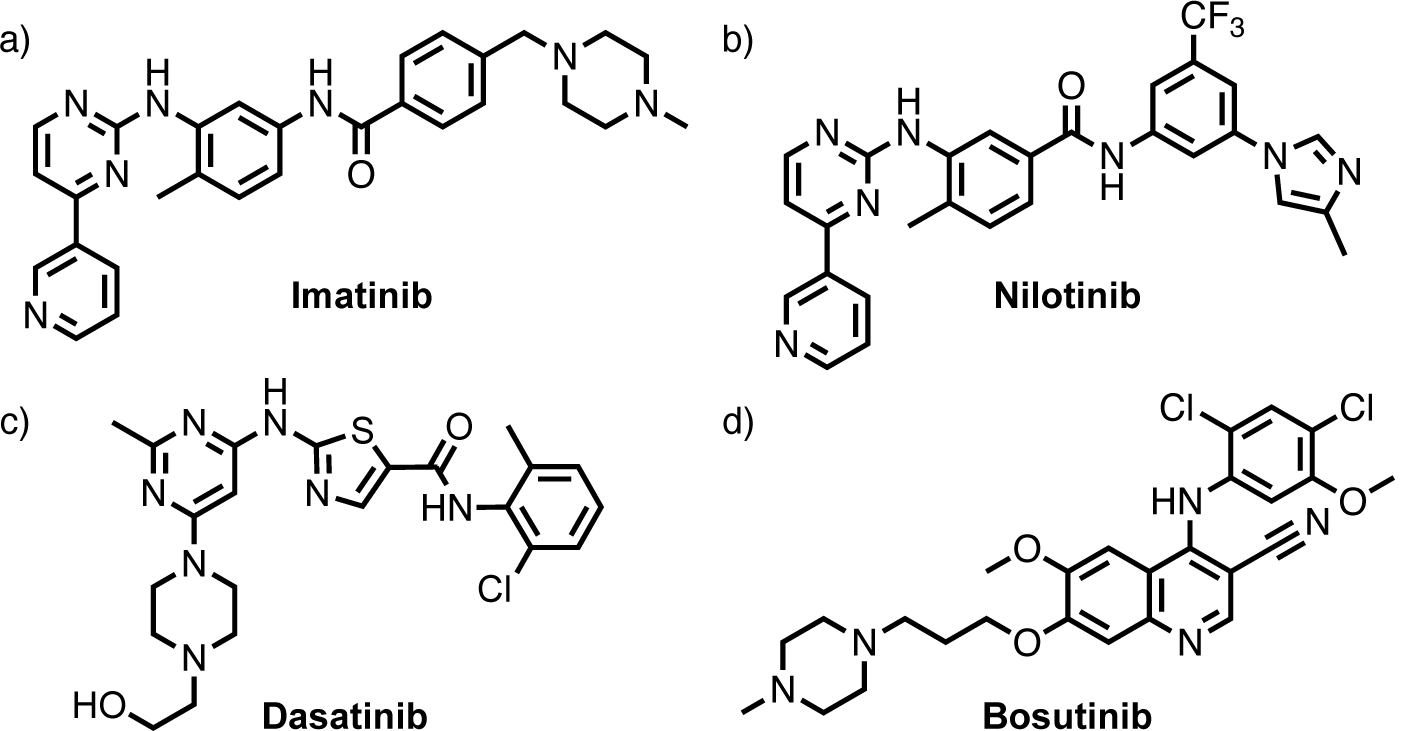
The four most common **TKI: imatinib** (a), **nilotinib**(b), **dasatinib** (c), and **bosutinib**
(d).

**Imatinib** is the first-choice therapy to treat the CML particularly in the early stages of the disease, also known as the chronic phase (CP). As patients reach the accelerated phase (AP) of CML they begin to exhibit no significant response to **imatinib**.^6^ This is due to the emergence of clones of CML with mutant forms of **BCR-ABL**.^1^ These mutations in **BCR-ABL** do not affect catalytic activity but decrease **imatinib** sensitivity. Disease resistance to **imatinib** has become a significant challenge in the treatment of CML.

The second-generation TKI are **bosutinib** (SKI-606), **dasatinib** (BMS-354825) and **nilotinib** (AMN-107), which all shows tremendous improvement in the treatment of the mutant forms of **BCR-ABL** (Scheme 1b-d). **Nilotinib** is structurally similar to that of **imatinib** and binds **BCR-ABL** similarly to that of **imatinib** with nearly 30 times higher affinity.^9^ **Bosutinib** and **dasatinib** can be classified as SRC kinase inhibitors as well as a **TKI. Dasatinib** can bind to both the active and inactive forms of **BCR-ABL** nearly 325 times more than **imatinib**.^10^ **Bosutinib** can interact with many **BCR-ABL** mutants and binds nearly 200 times more than **imatinib**.^10^ While second generation **TKI** show promise in treating CML in CP, there still is a need for more effective treatments for the AP of CML.

In order to help stimulate ideas for a 3^rd^ generation therapeutic, in this study we have designed two new novel drug candidates derived from **imatinib**. We have performed computational docking of these new candidates in **BCR-ABL** and compared how well these two new candidates are with **imatinib, dasatinib, bosutinib**, and **nilotinib**. Our two drug candidates have computationally higher docking scores than that of **imatinib**, and we predict that one candidate may be an orally available treatment. These candidates have the potential to be powerful new lead TKI.

## METHODS

The ATP competitive binding drug **imatinib** and the **BCR-ABL** kinase complexes were obtained from the RCSB website (1IEP).^11^ The crystal structure of **BCR-ABL** with **nilotinib** (3CS9)^12^, **bosutinib** (4MXX)^13^ and **dasatinib** (6BSD)^14^ inhibitors were used for computational modeling.

PyMol was used to visualize and measure interaction distances.^15^ ADMET properties were calculated using OpenEye FILTER, which screens drug candidates based on the bioavailability, toxicity, and the Lipinski rule of 5 (number of chiral centers, Lipinski hydrogen-bond donors, Lipinski hydrogen-bond acceptors, and molecular weight).^16^ VBrood is used to build new drug candidates via bioisosteric replacement using **imatinib** as the lead molecule.^17^

Gaussian09 was used to optimize molecules with AM1^18^ semi-empirical method.^19^

Conformation libraries were generated using OMEGA.^20^ Drug candidates were docked in the active site of BCR-ABL using FRED and FRED receptor.^21^ FRED and FRED receptor generates a total score, steric score, protein, ligand desolvation energies, and hydrogen bond scores. The steric score measures the steric contacts of heavy atoms between the ligand and enzyme interface. The protein and ligand desolvation terms are penalties from the loss of hydrogen bonds from implicit solvent molecules with the protein and ligand, respectively, when both groups are brought together in docking. The hydrogen bonds term is calculated from hydrogen bonds formed between the ligand and enzyme. Lastly, the total score is the sum of all of the energies (steric score, protein and ligand desolvation energies, and hydrogen bond scores). These methods have been previously utilized designing new dipeptidyl peptidase-4 inhibitors.^22^

## RESULTS AND DISCUSSION

### Evaluation of existing TKI

Currently the most popular drugs for the treatment of CML are **imatinib, nilotinib, dasatinib**, and **bosutinib**. We performed docking of each of the four drug molecules into the active site of the **BCR-ABL** (1IEP) using FRED and FRED receptor, and the results are tabulated in Table 1. All calculated chemgauss4 and ADMET values for each drug are tabulated in the supporting information (Table S1).

**Table 1.** FRED receptor and ADMET values from docking **imatinib, nilotinib, dasatinib**, and **bosutinib** with **BCR-ABL**.

|  | Total Score | Steric | Protein Desolvation | Ligand Desolvation | Hydrogen Bonds | LogP | Relative Binding Affinity |
| --- | --- | --- | --- | --- | --- | --- | --- |
| **Imatinib** | -10.47 | -14.28 | 3.74 | 4.68 | -4.62 | 2.59 | x1 |
| **Nilotinib** | -12.29 | -20.17 | 4.99 | 5.11 | -2.23 | 3.77 | x30 |
| **Bosutinib** | -9.41 | -17.00 | 4.27 | 5.04 | -1.72 | 3.64 | x200 |
| **Dasatinib** | -10.74 | -20.00 | 4.98 | 6.85 | -2.57 | 1.89 | x325 |

Among the four of the existing **BCR-ABL** inhibitors, the order of relative drug binding affinity is **imatinib, nilotinib, bosutinib**, and **dasatinib**. The computed value that most closely resembles the relative drug binding trend is the steric score, where the values of **dasatinib, nilotinib**, and **bosutinib** (–20.00, –20.17, and –17.00 respectively) are more favorable than the value of **imatinib** (– 14.28). The steric score represents how well these drugs fit in the **BCR-ABL** active site, and FRED correctly predicts that the more pharmaceutically effective drugs fit better in the **BCR-ABL** active site than **imatinib**, however FRED predicts that **dasatinib** and **nilotinib** are nearly equal in their predicted energy. The protein and ligand desolvation penalties inversely correlate with effectiveness, that being the larger the solvation penalty the more effective the drug. The hydrogen bond score does not correlate with drug effectiveness, this suggests that hydrogen bonding is not a significant interaction between the ligand and protein. In addition, the calculated LogP (cLogP) values would suggest not all of these drug molecules are orally available, which only **imatinib** and **dasatinib** can be taken orally.

These existing treatments of CML effectively interact with **BCR-ABL**, however we can gain a better understanding of how these drugs interact with **BCR-ABL**. Figure 2 displays **imatinib** redocked in **BCR-ABL** active site (Figure 2a) with key interactions highlighted between active site residues and **imatinib** (Figures 2b and 2c). Imatinib forms a linear shape within the active site cavity, coming into steric contact with most of the cavity residues. Hydrogen bonds are formed between residues E286, T315, and D381 with the ligand (Figure 2c). Moreover, the pyridine moiety of **imatinib**, which mimics ATP binding, forms a slip-stacked pi-pi interaction with F317 with distance 4.2Å (Figure 2c). We predict that these are key interactions that should be conserved in our drug designs and modifications from **imatinib**.

**Figure 2.**
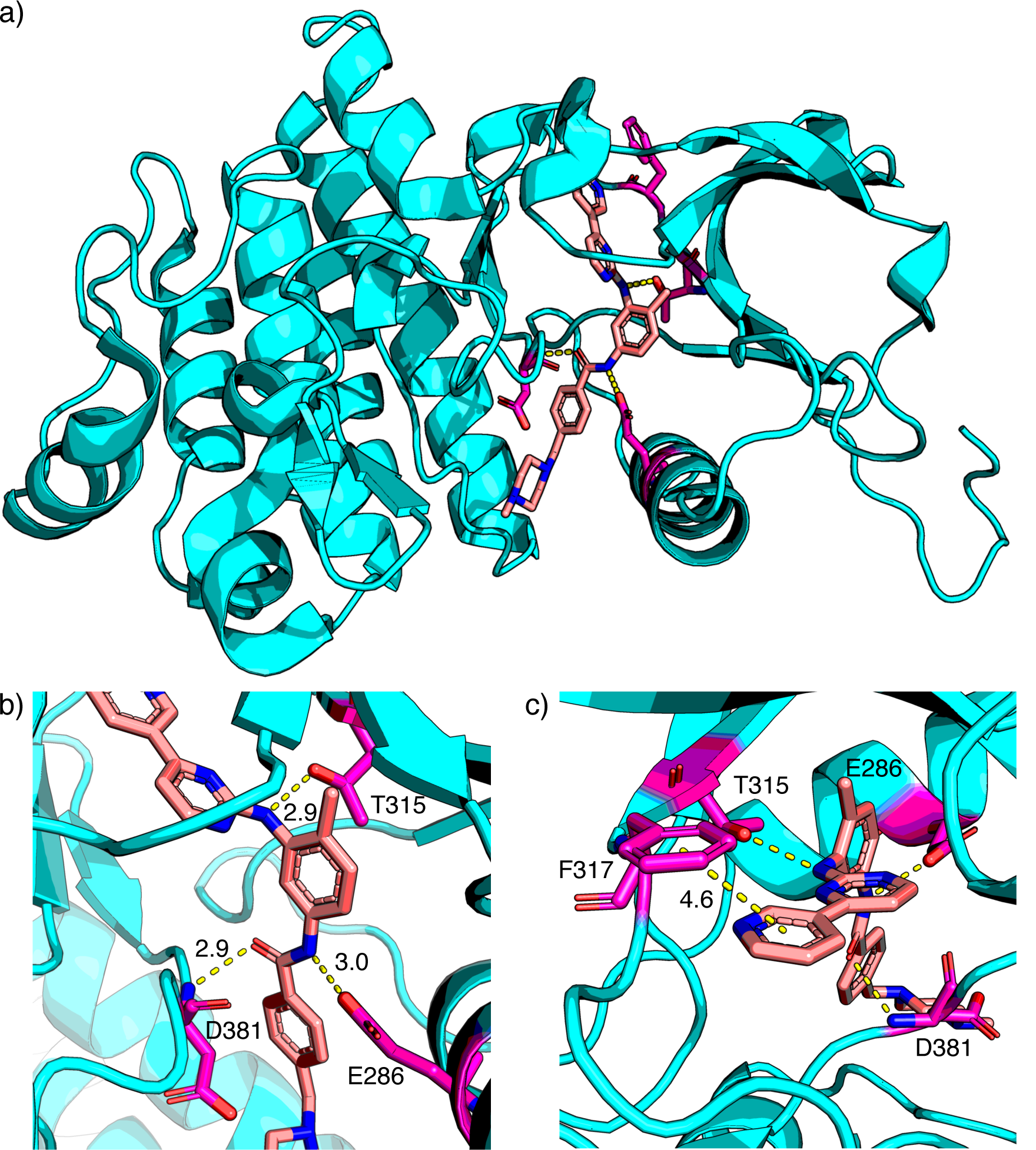
**Imatinib** (pink) redocked into **BCR-ABL** (teal) with interacting residues shown in purple (a). Residues E286, T315, and D381 are hydrogen bonding to **imatinib** (b). F317 forms a slip-stacked pi-pi stacking with the pyridine moiety of **imatinib** (c). Distances are reported in Ång-stroms (Å).

### Chemical Knowledge Driven Drug Design

We first designed a new drug candidate, **Candidate 1** (Figure 3), from **imatinib** based on pharmaceutical chemistry knowledge and structural analysis of existing drug molecules. We started our design with reconnecting the central amide moiety to be similar to that of **nilotinib**. We next replaced the piperazine group with an oxazole group and moved this group from the *meta* position to *para* to the conjoining amide. By changing the piperazine group to oxazole group, the potential toxicity of the molecule should be lowered by preventing the candidate from interfering with the metabolic pathways that involving the N-dealkylation of the piperazine ring.^23^ The oxygen atom of the oxazole may also promote an additional hydrogen bond within the active site. We also replaced the trifluoromethyl group to a fluoro group, which reduces the inductive effect to the benzene ring and also reduces the molecular weight. After several docking pre-trials, the fluoro and oxazole groups interact with **BCR-ABL** best at the *meta* and *para* positions, respectively.

**Figure 3.**
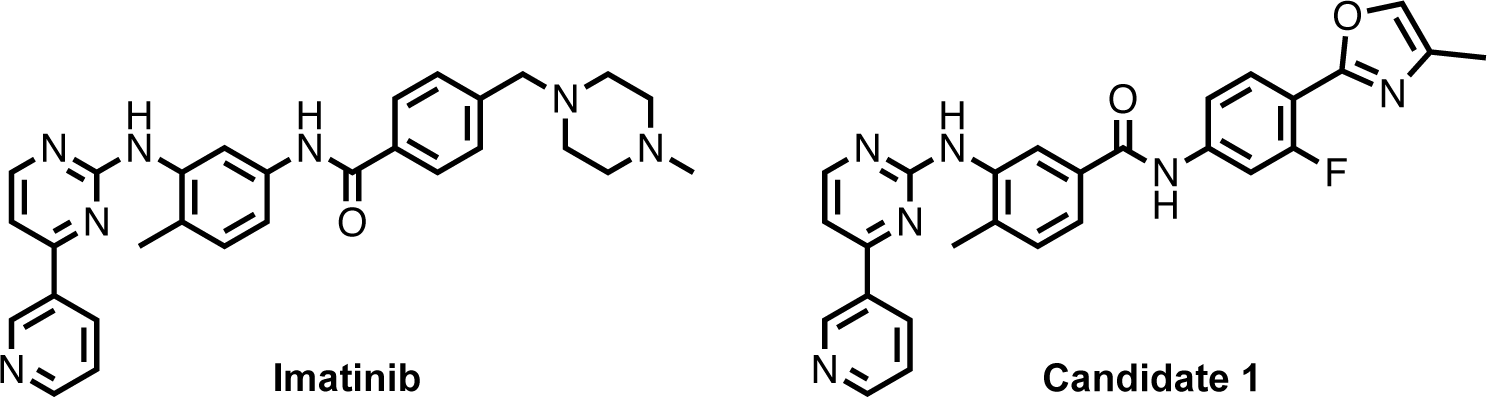
The structures of **imatinib** (left) and **Candidate 1** (right).

The computed docking values of **Candidate 1** with **BCR-ABL** are tabulated in table 2 with score comparisons to **nilotinib** and **imatinib**. Calculated chemgauss4 and ADMET values for **Candidate 1** are tabulated in the supporting information (Table S1). The docking score is more favorable for **Candidate 1** than **nilotinib** and **imatinib** by 0.44 and 2.26, respectively. The steric score for **Candidate 1** is lower than that of **imatinib** but higher than that of **nilotinib**, –1.26 and +4.63 respectively, which indicates that **Candidate 1** may be an effective TKI in the treatment of CML. Protein and ligand desolvation penalties are also less severe for **Candidate 1** than **nilotinib** and **imatinib**, and the hydrogen bond score is also more favorable. The calculated LogP value is similar to **bosutinib** and **nilotinib**, which we predict that **Candidate 1** might be less orally available than **dasatinib** and **imatinib**.

**Table 2.** Computed docking values of Candidate **1** with comparisons to **imatinib** and **nilotinib**.

|  | Total Score | Steric | Protein Desolvation | Ligand Desolvation | Hydrogen Bonds | LogP |
| --- | --- | --- | --- | --- | --- | --- |
| **Candidate 1** | -12.73 | -15.54 | 3.02 | 4.76 | -4.96 | 3.50 |
| Δ**Nilotinib** | -0.44 | +4.63 | -1.97 | -0.59 | -2.73 | -0.27 |
| Δ**Imatinib** | -2.26 | -1.26 | -0.72 | -0.08 | -0.34 | +0.91 |

The computed properties from docking **Candidate 1** with **BCR-ABL** leads us to believe we potentially have a new lead molecule for future CML treatment. To better understand how this drug candidate interacts with **BCR-ABL**, we investigated the molecular interactions of **Candidate 1** with **BCR-ABL**, which is shown in Figure 4. Unlike **imatinib, Candidate 1** is predicted to bend into a “U” shape within the active site cavity, which limits the steric contacts that made in the active site cavity (Figure 4a), which likely contributes to the lower steric score. The pyridine moiety of **Candidate 1** is facing the “southern” side of the active site and forming a hydrogen bond with the D381 carbonyl backbone, rather than pi-pi stacking interaction with F317 (Figure 4b). The new oxazole moiety forms a new hydrogen bond with K271 (Figure 4b). Candidate 1 does not interact with E286, T315, and F317 (Figure 4c). The methyl group and the double bond contribute to the lipophilic interactions to further stabilize the structure in the active site. **Candidate 1** also interacts with the activation loop, similarly to **imatinib** (Figure 4d).

**Figure 4.**
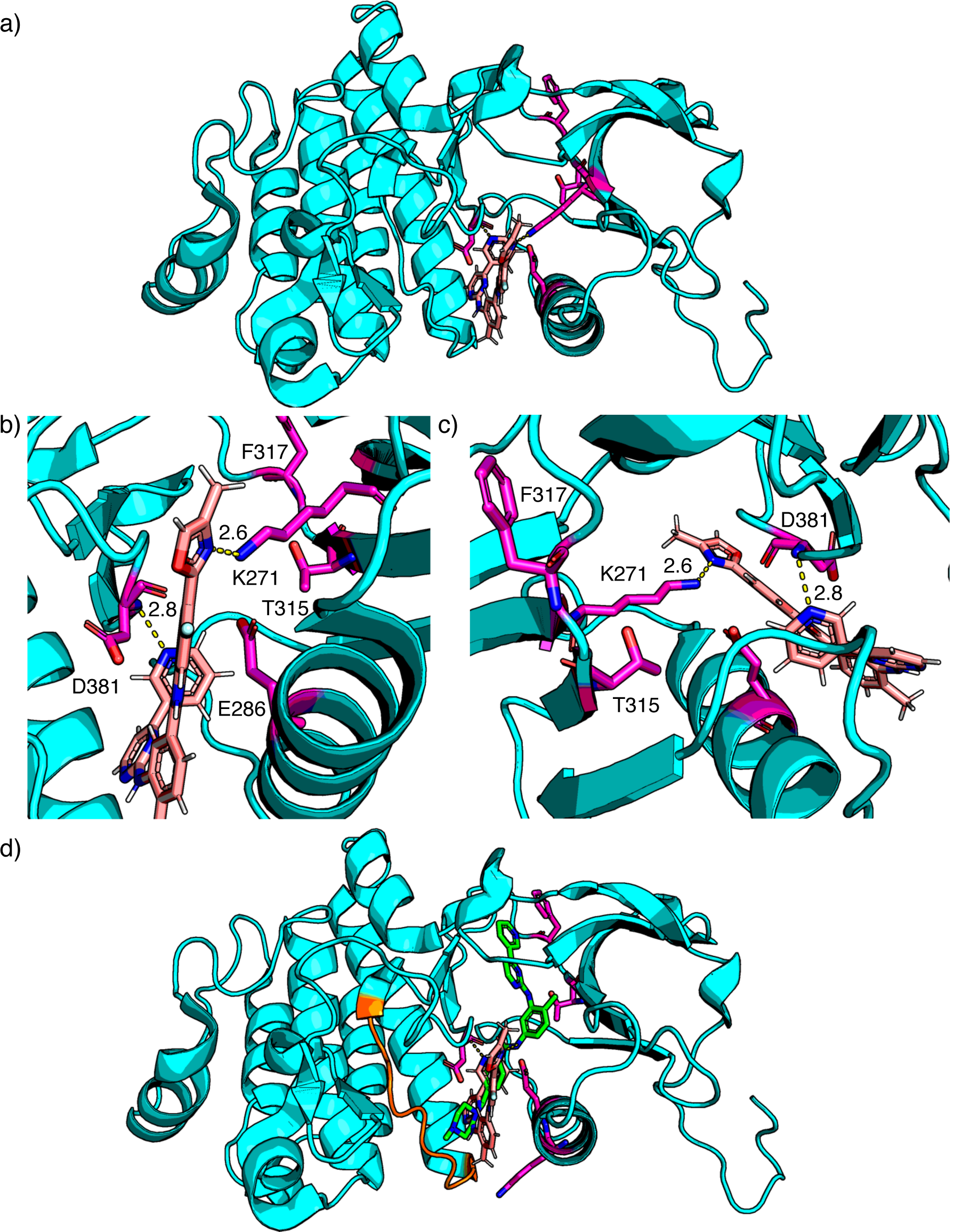
**Candidate 1** (pink) docked into **BCR-ABL** (teal) with interacting residues shown in purple (a). Residues K271 and D381 are hydrogen bonding to **Candidate 1** (b & c). Overlay of **Candidate 1** (pink) and **imatinib** (green) docked into **BCR-ABL** and interacting with the activation loop (orange) (d). Distances are reported in Ångstroms (Å).

In summary, the chemical knowledge and intuition drug designed TKI **Candidate 1** displays an overall desirable result compared to **nilotinib** and **imatinib**. The total score is the lowest among all four drugs, and the steric score is lower than **imatinib**, which suggests **Candidate 1** could be a new lead molecule.

### Computationally Driven Drug Design

While **Candidate 1** was designed starting from **imatinib** using chemical knowledge and intuition, we also explored the design of a new drug candidate (**Candidate 2**) using the computational software VBrood^17^ (Figure 5). VBrood identified that the piperazine moiety to be replaced by a cationic *n*-methyldihydro-imidzole and that the central amide be reconnected similar to that of **nilotinib**.

**Figure 5.**
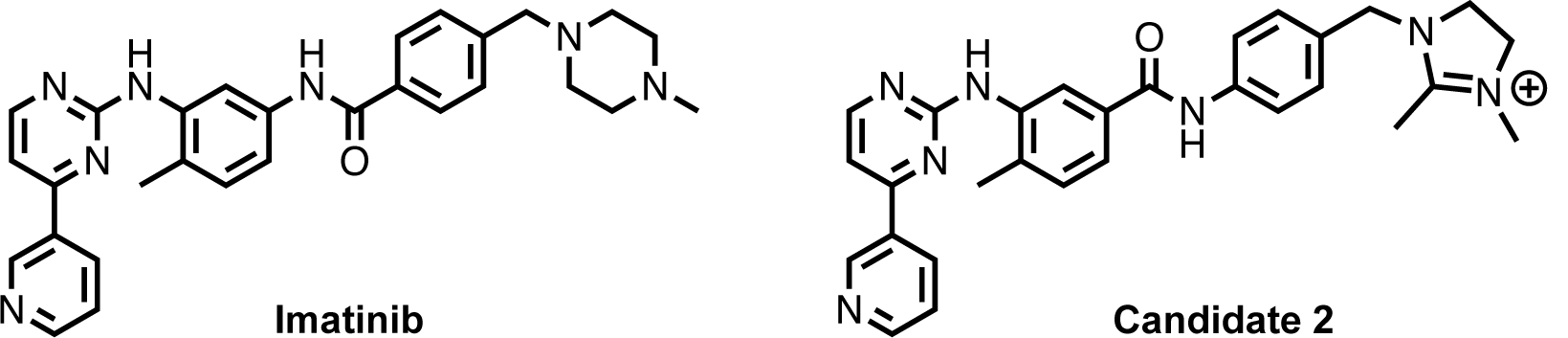
The structures of **imatinib** (left) and **Candidate 2** (right).

The docking results for **Candidate 2** are listed in table 3. Calculated chemgauss4 and ADMET values for **Candidate 2** are tabulated in the supporting information (Table S1). **Candidate 2** has the lowest total score of –13.95 among all of the drugs and **Candidate 1**. The steric score for **Candidate 2** is lower than **imatinib** and **Candidate 1** but higher than **nilotinib**, suggesting that this candidate might be a promising new lead molecule. The hydrogen bond score is also lower than **imatinib, nilotinib**, and **Candidate 1**. The LogP value of **Candidate 2** is lower than **nilotinib** and slightly higher than **imatinib**, suggesting that this drug candidate is likely equally as orally available as **dasatinib** and **imatinib**. Based on all of the computed scores we predict that the computationally designed drug (**Candidate 2**) would perform the best compared to the chemical knowledge-based designed drug (**Candidate 1**) and the four common treatments.

**Table 3.** Computed docking values of **Candidate 2** with comparisons to **imatinib** and **nilotinib**.

|  | Total<br>Score | Steric | Protein<br>Desolvation | Ligand<br>Desolvation | Hydrogen<br>Bonds | LogP |
| --- | --- | --- | --- | --- | --- | --- |
| **Candidate 2** | -13.95 | -16.93 | 4.40 | 4.25 | -5.68 | 2.68 |
| **ΔNilotinib** | -1.66 | +3.24 | -0.59 | -0.86 | -3.45 | -1.09 |
| **ΔImatinib** | -3.48 | -2.65 | +0.66 | -0.43 | -1.06 | +0.09 |

The key interactions between residues in the active site and **Candidate 2** are displayed in Figure 6. Similar to **imatinib, Candidate 2** forms a linear shape within the active site cavity. Yet the pyridine moiety faces the “southern” side of the cavity similar to that of **Candidate 1** (Figure 6a). **Candidate 2** also hydrogen bonds to D381 and E286, and K285 hydrogen bonds to pyridine moiety (Figure 6b). The hydrogen bond to D381 is shorter by 0.4 Å respective to **imatinib**. The cationic imidazole moiety is within the region of T315 and F317. However, this group does not interact with these residues (Figure 6c). **Candidate 2** also interacts with the activation loop, similarly to **imatinib** (Figure 6d).

**Figure 6.**
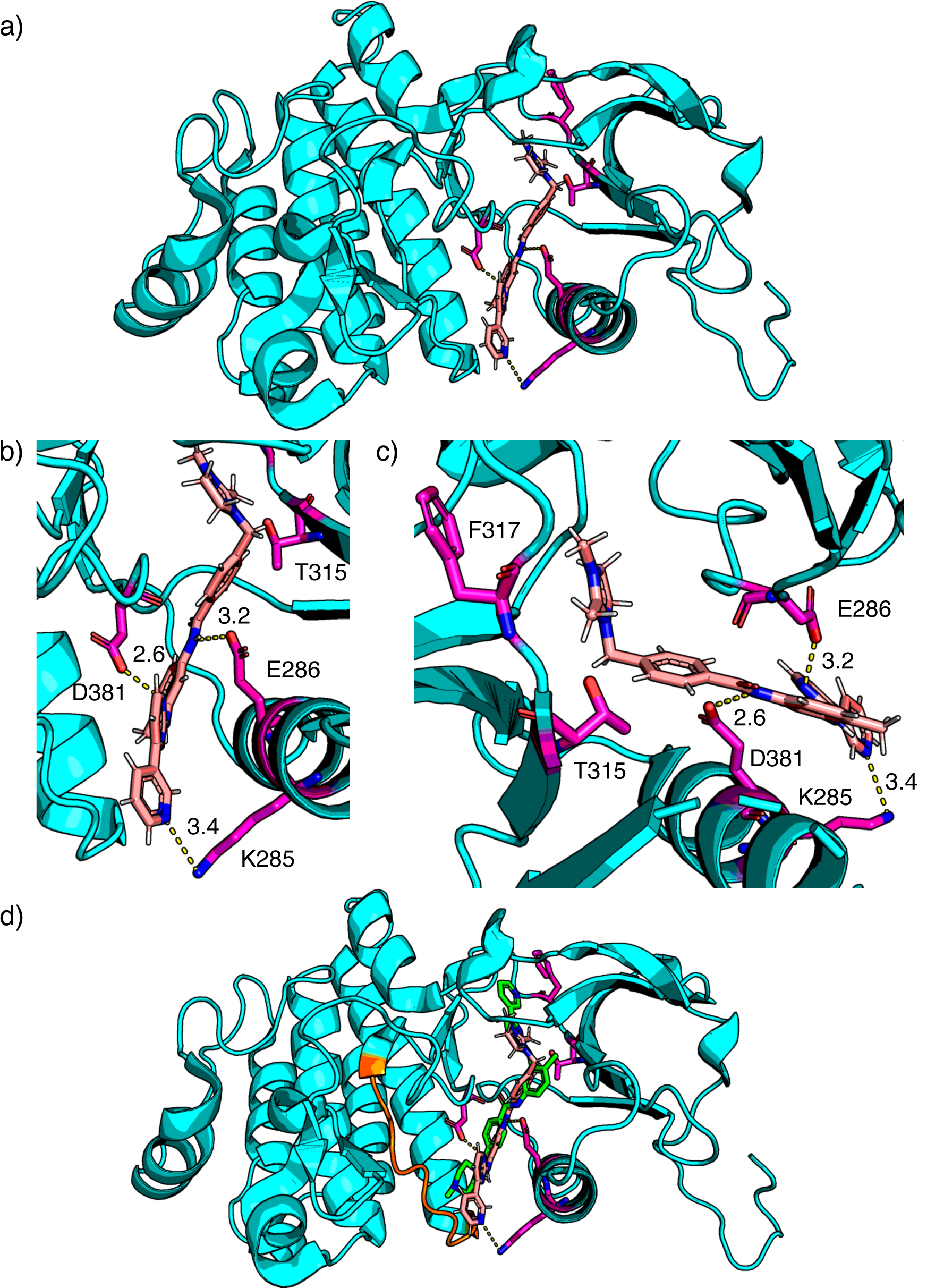
**Candidate 2** (pink) docked into **BCR-ABL** (teal) with interacting residues shown in purple (a). Residues K285, E286, and D381 are hydrogen bonding to **Candidate 1** (b & c). Overlay of **Candidate 2** (pink) and **imatinib** (green) docked into **BCR-ABL** and interacting with the activation loop (orange) (d). Distances are reported in Ångstroms (Å).

## CONCLUSION

As CML becomes resistant to the first line treatment **imatinib**, second generation drugs, **nilotinib, dasatinib**, and **bosutinib**, are utilized to inhibit **BCR-ABL** as the disease progresses to the CP. New TKI are needed in order to bypass drug-resistant mutants of **BCR-ABL**.^24^ We generated two new drug candidates by performing modifications on **imatinib** using chemical knowledge and intuition (**Candidate 1**), and using computational software for bioisosteres replacement (**Candidate 2**). Both of the drug candidates generated are predicted to be more effective than **imatinib** as well as potentially **nilotinib**. Even though these drug leads are predicted to be more advanced treatments to CML, further testing and studies *in vivo* are required to determine true efficacy. Despite the further studies that need to be conducted in the future, the usage of computational software and chemical knowledge serve as a good starting point to propose new drug molecules for advanced treatments.^25^

## AUTHOR INFORMATION

### Notes

The authors declare no competing financial interest.

