## Supporting Information for "Computationally Guided Design of BCR-ABL Tyrosine Kinase Inhibitors"

Chenghao Duan[1], Simon K. S. Chu[2], Justin B. Siegel[1,3,4] and Jason S. Fell[*,4]
[1]Department of Chemistry, University of California Davis, Davis, California, United States of America
[2]Biophysics Graduate Group, University of California, Davis, California, United States of America
[3]Department of Biochemistry & Molecular Medicine, University of California, Davis, California, United States of America
[4]Genome Center, University of California, Davis, California, United States of America

**Table S1.** Chemgauss4 and ADMET values from docking **imatinib**, **nilotinib**, **dasatinib**, **bosutinib**, **Candidate1**, and **Candidate 2** with **BCR-ABL**.

| | Total Score | Steric | Protein Desolvation | Ligand Desolvation | Hydrogen Bonds | LogP | Chiral Centers | H-Bond Donors | H-Bond Acceptors | Molecular Weight |
|---|---|---|---|---|---|---|---|---|---|---|
| **Imatinib** | −10.47 | −14.28 | 3.74 | 4.68 | −4.62 | 2.59 | 0 | 2 | 8 | 494.61 |
| **Nilotinib** | −12.29 | −20.17 | 4.99 | 5.11 | −2.23 | 3.77 | 0 | 2 | 8 | 529.52 |
| **Dasatinib** | −10.74 | −20.00 | 4.98 | 6.85 | −2.57 | 1.89 | 0 | 4 | 9 | 489.01 |
| **Bosutinib** | −9.41 | −17.00 | 4.27 | 5.04 | −1.72 | 3.64 | 0 | 2 | 8 | 531.45 |
| **Candidate 1** | −12.73 | −15.54 | 3.02 | 4.76 | −4.96 | 3.50 | 0 | 2 | 8 | 480.49 |
| **Candidate 2** | −13.95 | −16.93 | 4.40 | 4.25 | −5.68 | 2.68 | 0 | 2 | 8 | 492.59 |
